# DEEP: Extracting Latent Energetic Landscapes from Protein Structure Predictors

**DOI:** 10.64898/2026.10.15.759389

**Authors:** Zsófia Rózsa, Gábor Erdős

## Abstract

Modern protein structure predictors learn complex sequence-structure relationships, yet the extent to which their internal representations capture the energetic principles governing protein stability and conformational preferences remains unclear. Here, we introduce DEEP (Distogram Encoded Energetic Potential), a general framework that extracts effective statistical potentials directly from predicted residue-pair distance distributions, using the model’s own terminal distance state as an internal reference without training on experimental stability data. Applied to Boltz-2 (DEEP-Boltz) achieves a mean per-assay Spearman correlation of 0.720 across 64 ProteinGym stability assays, numerically outperforming all 97 published zero-shot methods in the matched comparison on par with the best performing predictors trained directly on experimental stability measurements. Beyond mutation effects, DEEP-Boltz accurately distinguishes native-like from non-native conformations across more than 180,000 Rosetta decoys. Applying DEEP to AlphaFold2 reveals nearly identical conformational discrimination but substantially weaker mutation-effect prediction, demonstrating that recognizing favorable conformations and capturing their energetic sensitivity to sequence changes are distinct properties of learned structural representations. Together, these findings establish that protein structure predictors encode accessible energetic information beyond their geometric predictions and that this information can be extracted. We release the framework and DEEP-Boltz as a freely available tool for extracting these latent energetic representations and for zero-shot prediction of mutation-induced protein stability changes.

## 1. Introduction

The emergence of highly accurate protein structure predictors has transformed the relationship between protein sequence and three dimensional structure. Modern models such as AlphaFold2 and Boltz learn complex sequence dependent constraints that allow them to identify geometries compatible with a protein sequence with remarkable accuracy ^1-3^. Their outputs, however, are predominantly interpreted geometrically, as predicted coordinates, residue pair distances, or measures of structural confidence. A more fundamental question is what these models have learned about the physical organization underlying protein structure. Distinguishing compatible from incompatible conformations requires learning strong constraints on residue packing and long range organization, raising the possibility that representations trained for structure prediction also contain information related to the energetic landscape of proteins ^4-7^.

Such information would provide a largely unexplored connection between modern structure prediction and classical statistical descriptions of protein energetics. Statistical potentials infer effective interactions from the relative occurrence of structural states, exploiting the statistical mechanical relationship between probability and free energy ^8^. Traditionally, these potentials are constructed explicitly from databases of experimentally determined protein structures and require a reference distribution against which observed structural frequencies are compared ^9^. Structure predictors now learn analogous relationships implicitly ^10^. Given a sequence, they assign preferences to alternative structural configurations based on large collections of structural and evolutionary information ^11^. Whether these learned preferences can be transformed into an explicit and experimentally useful energetic representation, however, remains unclear.

Residue pair distance distributions (distograms) provide a natural representation in which to investigate this question ^12^. Rather than specifying only a single predicted structure, a distogram assigns a sequence conditioned probability distribution to the possible separations of each residue pair. These distributions therefore describe not only the geometry ultimately selected by the predictor, but also its relative support for alternative geometric states ^13^. In principle, this information resembles the probability distributions from which statistical potentials are derived. A high probability for a particular distance, however, cannot itself be interpreted as an interaction energy ^14^. The probability must be evaluated relative to an appropriate reference state, and defining this reference is one of the central problems in constructing knowledge based potentials ^15^.

Classical statistical potentials address this problem using externally defined reference distributions, such as ideal gas states, randomized structures, or empirical backgrounds derived from structural databases ^16^. Learned structure predictors present a different opportunity. Their distance representations can contain both probabilities for specific folded geometries and an open ended state corresponding to residue pairs being spatially separated. This raises the possibility that the predictor itself may provide the reference required to interpret its geometric probabilities energetically. An observed residue pair geometry could then be evaluated relative to loss of that specific interaction through spatial separation, producing a model internal statistical potential without constructing an external background distribution or training an additional energetic model.

A stringent test of such a representation is whether it captures experimentally measured changes in protein stability. Folding stability reflects the free energy difference between folded and unfolded ensembles, and a sequence substitution changes this balance by ΔΔ*G* =Δ*G*(*s*_mut_) − Δ*G*(*s*_WT_)^17^. Predicting these changes is challenging because mutations can simultaneously perturb packing, hydrogen bonding, electrostatics, solvation and conformational entropy, with their consequences depending strongly on the local three dimensional environment ^18^. Protein stability therefore provides a direct experimental setting in which to ask whether the sequence response of a learned structural representation contains physically meaningful energetic information.

A wide range of computational approaches have been developed for this problem ^19,20^. Physics based methods estimate mutation effects using molecular mechanics, empirical energy functions or molecular simulation, while supervised predictors learn relationships between structural or sequence features and experimentally measured stability changes ^21-24^. More recently, protein language models, inverse folding models and structure aware sequence representations have enabled powerful zero shot mutation predictions without direct training on stability measurements ^25-28^. Large scale datasets such as the Tsuboyama proteolysis experiments, incorporated into the ProteinGym Stability or the S669 stability benchmark set, now provide systematic measurements across large numbers of variants and allow such models to be compared under a common framework ^29,30^. These data therefore offer an unusually stringent test of whether an energetic quantity extracted directly from a structure predictor contains experimentally relevant information.

Importantly, mutation prediction probes only one axis of an energetic landscape with the sequence changing while the structural state is held fixed. A complementary test is obtained by holding the sequence fixed while varying structure. An effective potential that reflects the conformational preferences learned by a structure predictor should assign more favorable scores to native like than to increasingly distorted conformations ^9,31^. Considering these perturbations together distinguishes two properties that are often treated implicitly as equivalent: recognizing conformations compatible with a sequence and responding energetically when that sequence is perturbed. Comparing these properties across independently developed structure predictors can therefore reveal how learned structural representations organize energetic information.

Here we introduce DEEP (Distogram Encoded Energetic Potential), a general framework for extracting an effective statistical potential directly from the sequence conditioned distance distributions of frozen protein structure predictors. DEEP compares the probability density assigned to an observed residue pair geometry with the predictor’s own terminal far distance state, using the latter as a model internal representation of spatial separation. Because the potential is obtained directly from the predictor output, no experimental energetic measurements are required to construct or train the model.

We first implement the framework using Boltz-2 (DEEP-Boltz) and test whether mutation induced changes in the extracted potential correspond to experimentally measured stability changes. DEEP-Boltz reaches a mean per-assay Spearman correlation of 0.720 across 64 matched ProteinGym Stability assays, numerically exceeding all 97 published zero-shot predictors in the matched comparison and approaching the performance of more recent methods trained directly on experimental stability data. The mutation sensitive signal is strongly localized around the substituted residue, with a broad optimum at approximately 6-8 Å. We then perturb the complementary axis of the landscape and show that DEEP strongly discriminates native like from non native structures across large conformational decoy ensembles.

Finally, we implement the same framework using AlphaFold2 (DEEP-AF2) to determine whether these properties generalize across structure predictors. Boltz-2 and AlphaFold2 produce nearly identical rankings of alternative conformations but differ substantially in their response to sequence substitutions. Thus, learning the conformational landscape and learning its energetic sensitivity to sequence are related but separable properties of a structure predictor. Together, these results show that sequence conditioned distograms contain an accessible latent energetic representation that extends beyond their conventional interpretation as intermediate geometric predictions. DEEP provides a general route for extracting this information, allowing frozen structure predictors to be interrogated as energetic models rather than coordinate generators alone.

## 2. Results

Our central hypothesis is that structure predictor distograms contain an energetic representation that can be recovered without retraining the underlying model. We first define DEEP and test its relationship to experimentally measured mutation-induced stability changes, then examine whether the same potential captures conformational preferences and whether these properties generalize across independently developed structure predictors.

### 2.1 DEEP converts sequence conditioned distograms into an effective statistical potential

DEEP starts from the pairwise distance distributions produced internally by a protein structure predictor. Rather than using only the final predicted coordinates, we ask whether these distributions can be transformed into an energy-like representation of the geometries favored by a given sequence. The construction itself has three components: correcting distance probabilities for configurational volume, defining a model internal reference state for each residue pair, and integrating these pairwise preferences into a potential for a complete structure.

For a sequence *s* and residue pair *i,j*, the structure predictor assigns a probability *p*_*ij*_(*b*∣*s*) to each distance bin *b* (Figure 1A). These probabilities cannot be compared directly across distances because bins at larger radii represent larger configurational volumes. We therefore convert the probability assigned to each bin into a radial log density *l*_*ij*_,

**Figure 1:**
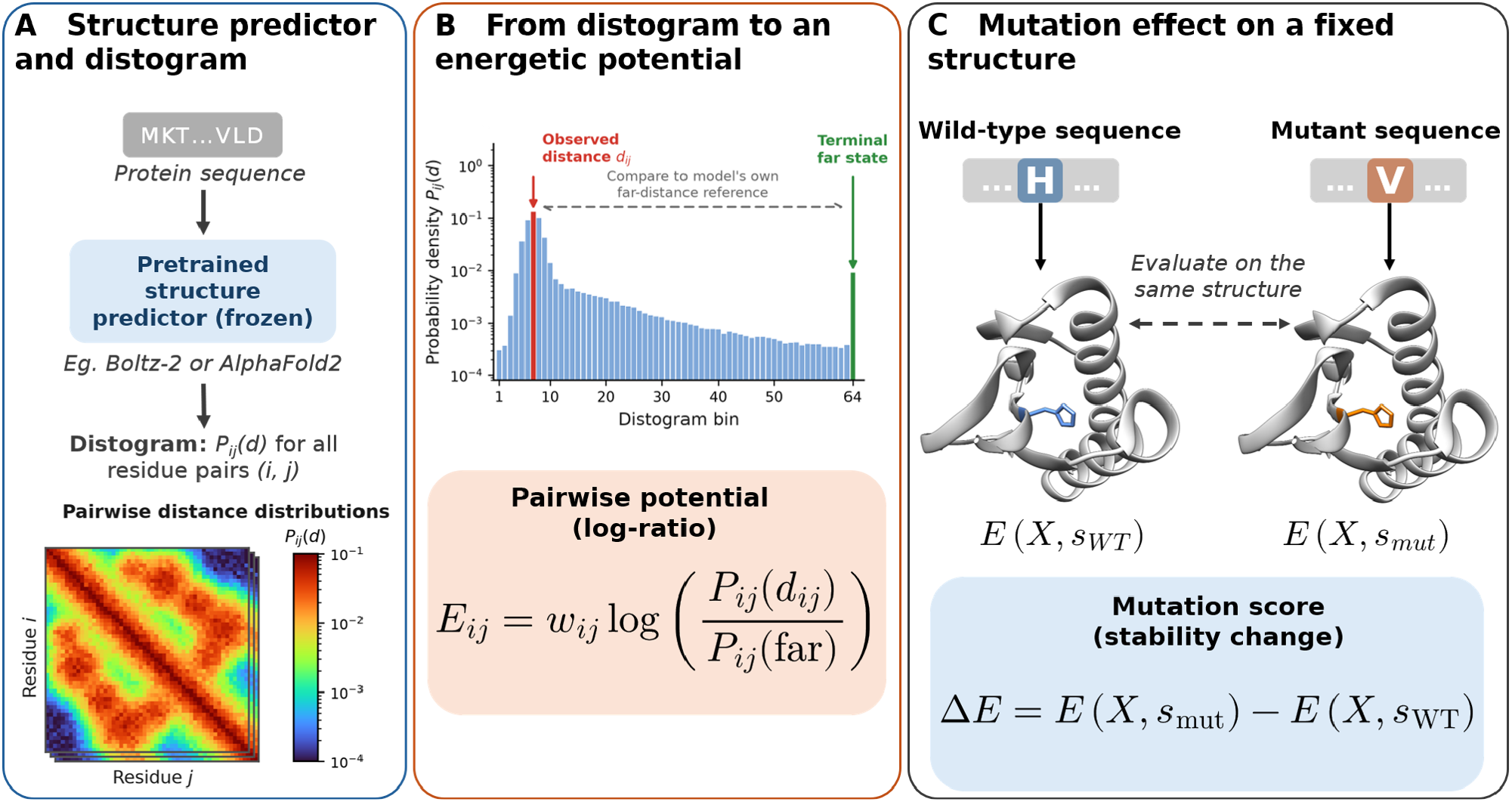
General overview of the DEEP framework. **(A)** A pretrained, frozen structure predictor generates sequence-conditioned residue-pair distance distributions (distograms). **(B)** DEEP converts these distributions into an effective pairwise statistical potential by comparing the probability of an observed residue-pair distance with that of the terminal far-distance state, which serves as a model-internal reference. **(C)** Mutation effects are evaluated by scoring wild-type and mutant sequences against the same fixed structure and calculating the difference in their effective potentials.

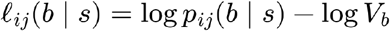

where *V*_*b*_ is the spherical shell volume represented by bin *b*. This correction separates enrichment at a particular distance from the purely geometric increase in the number of configurations available at larger radii. The resulting quantity can be interpreted as an effective statistical preference of the model for a residue pair to occupy that distance.

A statistical potential does not only require a probability for the observed state but also a reference against which that probability is evaluated. Classical statistical potentials obtain this reference from an externally specified background distribution ^8^. DEEP instead derives it from the structure predictors internal representation (Figure 1B). For each residue pair, we compare the density assigned to its observed distance *d*_*ij*_ in the folded structure with that assigned to the model’s terminal distance state *b*_far_:

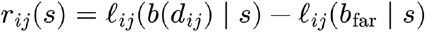

This log ratio expresses how strongly the predictor favors the observed folded residue pair geometry over spatial separation for this particular sequence. The terminal state therefore provides a sequence and pair specific internal reference rather than a universal background imposed externally. We use it operationally as a model internal separated state, analogous to the loss of a specific folded pair interaction, without assuming that it represents the complete thermodynamic unfolded ensemble.

The pairwise terms can then be combined to obtain an effective potential for a complete structure *X*:

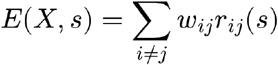

The resulting quantity is a model-internal effective potential *E* assigned to structure *X* given sequence . It is not assumed to be a thermodynamic free energy on an absolute scale, instead it measures the compatibility of a particular sequence-structure combination relative to the predictor’s own separated-state reference. Because the potential depends explicitly on both *X* and, either variable can subsequently be perturbed while the other is held fixed. Here, *w*_*ij*_ determines how strongly each residue pair contributes to the total score. For mutation prediction, weights are calculated from the wild type prediction and subsequently held fixed. We use the entropy of the wild type distogram together with the probability assigned to the observed distance,

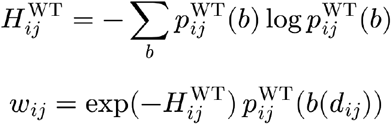

so that residue pairs for which the predictor gives a concentrated distribution and strong support to the observed geometry contribute more strongly. Importantly, these weights are a confidence based weighting of the learned pair representation rather than a thermodynamic entropy term.

To predict the effect of a sequence substitution, wild type and mutant sequences are evaluated against the same wild type structure and with the same pair weights (Figure 1C). The mutation score is therefore defined as:

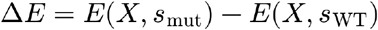

Holding the reference backbone fixed allows DEEP to assess how sequence substitutions alter the predictor’s support for the existing folded geometry without generating separate structures for each mutant. Substitutions between non-glycine residues preserve the occupied distance bins, whereas substitutions involving glycine require reassignment because the residue-centre convention changes from Cβ to Cα. Mutation effects are calculated from changes in the model-derived pairwise log-probability ratios without radial volume correction. The latter is applied only to conformational scoring, where alternative structures occupy different residue-pair geometries.

Finally, mutation induced energetic perturbations are expected to be concentrated around the substituted residue rather than distributed uniformly throughout the protein. DEEP therefore allows the pairwise sum to be restricted to residue pairs whose structural environment lies within a radius *R* of the mutation. We use *R* = 6 Å as the primary formulation, corresponding approximately to the immediate local interaction environment.

Together, these steps convert a structure predictor’s distogram into an interpretable mutation score. The model’s pairwise geometric probabilities are converted into statistical preferences, referenced to an internal separated state, integrated over the relevant structural environment, and expressed as a difference between wild type and mutant sequences. No experimental stability or other energetic measurements are used in this construction, thus DEEP can be considered as a complete zero shot predictor.

### 2.2 DEEP-Boltz reaches leading zero shot accuracy

We first implemented DEEP using the sequence-conditioned distograms of Boltz-2, which we refer to as DEEP-Boltz. Boltz-2 provides a 64-bin residue-pair distance distribution spanning 2-22 Å, with the terminal open-ended bin serving as the model-internal separated-state reference defined above. A systematic ablation of 67 alternative reference definitions supported this choice, with the terminal distance state producing the strongest mutation-effect correlation for DEEP-Boltz (Supplementary Note S1).

We evaluated DEEP-Boltz on the ProteinGym Stability benchmark derived from the Tsuboyama proteolysis dataset ^29^. The matched comparison contains 64 assays and 117,811 experimentally characterized mutations, and published ProteinGym predictions were re-aggregated over the same assay set. Using the local 6 Å formulation introduced above, DEEP-Boltz reaches a mean per-assay Spearman correlation of 0.720 across all mutations and 0.726 for single substitutions. This is numerically the highest mean correlation among the 97 published methods evaluated on the matched ProteinGym assays (Figure 2). Since the release of ProteinGym, several additional methods have been developed using its stability data for supervised training. Among these, ThermoMPNN reaches a correlation of 0.725, only marginally higher than the zero-shot performance of DEEP-Boltz ^24^. Thus, an energetic potential extracted directly from a frozen structure predictor approaches the performance of methods explicitly optimized on experimental stability measurements.

**Figure 2:**
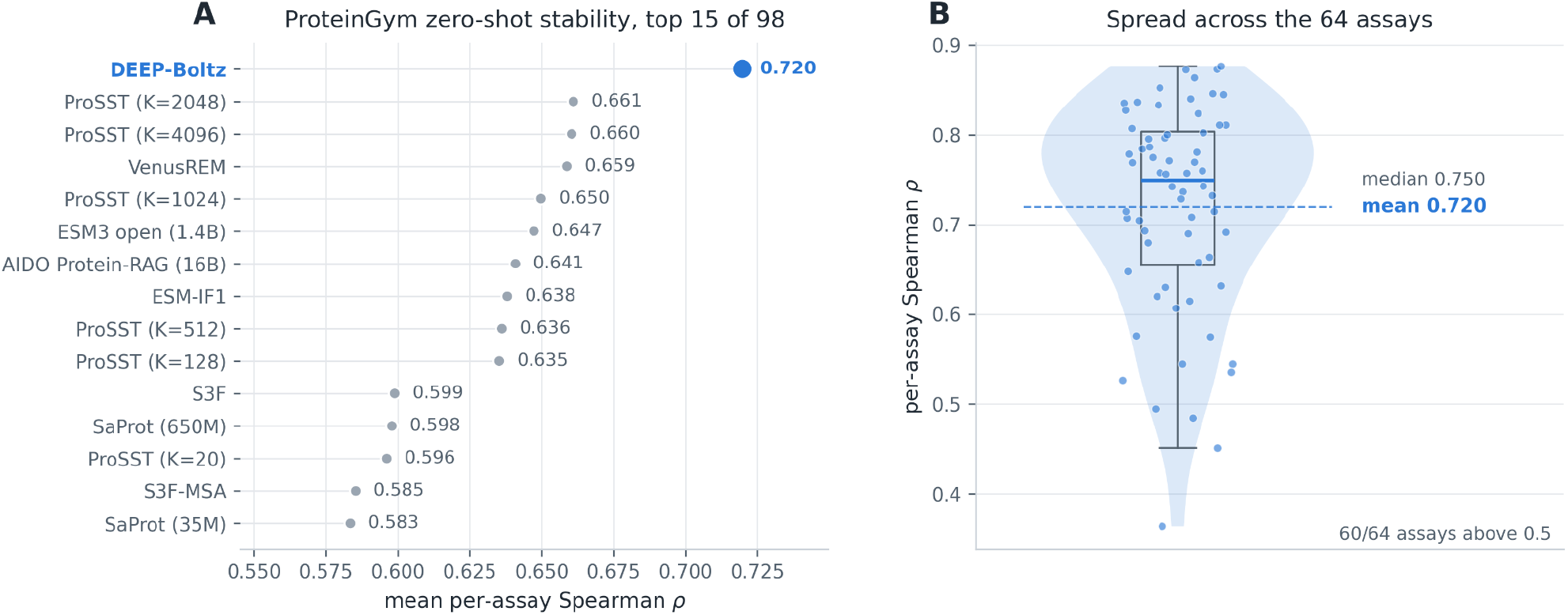
DEEP-Boltz achieves leading zero-shot mutation-stability prediction on ProteinGym. **(A)** Mean per-assay Spearman correlation across the 64 matched ProteinGym Stability assays for DEEP-Boltz and the top-performing published zero-shot methods. Using the 6 Å local formulation, DEEP-Boltz reaches a mean correlation of 0.720, numerically exceeding all 97 published methods evaluated on the same assay set. **(B)** Distribution of DEEP-Boltz performance across the 64 individual assays. Each point represents one assay; the violin and box plot show the overall distribution, with a mean Spearman correlation of 0.720 and a median of 0.750. DEEP-Boltz achieves a positive correlation exceeding 0.5 in 60 of 64 assays.

We next examined whether DEEP-Boltz depends on experimentally determined protein structures or can achieve comparable accuracy using structures predicted by Boltz-2 itself. Replacing experimental wild-type structures with Boltz-2 predictions resulted in essentially unchanged mutation-effect prediction performance, with comparable correlations across the evaluated assays. Thus, the mutation-sensitive energetic information extracted by DEEP-Boltz does not require experimentally determined structural coordinates, enabling its application to proteins for which only amino-acid sequences are available.

We next tested whether this predictive signal transfers beyond the ProteinGym benchmark. On the independent S669 dataset ^32^, which aggregates experimental stability measurements from 93 publications, DEEP-Boltz reaches a mean within-protein Spearman correlation of 0.413. Despite the substantially greater experimental heterogeneity of S669, this places DEEP-Boltz third among the 21 methods in the matched within-protein comparison and first among the zero-shot methods (Supplementary Note 2). Together, the ProteinGym and S669 results show that DEEP extracts mutation-sensitive information from Boltz-2 that is quantitatively related to experimental protein stability without training on stability measurements.

The pairwise formulation of DEEP additionally allows us to ask where this mutation-sensitive information is encoded. Rather than treating the 6 Å neighbourhood simply as a scoring choice, we systematically expanded the structural region contributing to each mutation score, from the immediate environment of the substituted residue to the complete protein.

### 2.3 Mutation sensitive energetic information is spatially localized

The predictive signal was strongly concentrated around the site of mutation. Because DEEP is an explicit sum over residue pairs, its mutation score can be spatially decomposed without modifying the underlying potential. For a mutation at position *p*, we define the cumulative score within structural radius *R* as:

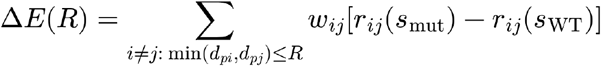

Increasing the radius revealed a pronounced but broad local optimum. Across the 64 ProteinGym assays, mean per-assay Spearman correlation rises from 0.668 at 4 Å to 0.720 at 6 Å and remains essentially level at 8 Å (0.718), before progressively declining as more distant residue pairs are included. The complete, untruncated potential reaches only 0.657. Single substitutions show the same profile, increasing from 0.643 at 4 Å to 0.726 at 6 Å and peaking at 0.731 at 8 Å before declining to 0.661 for the complete sum (Figure 3). So the most informative response occurs within approximately the first two interaction shells surrounding the substituted residue, whereas progressively incorporating distant responses reduces mutation-ranking accuracy.

**Figure 3:**
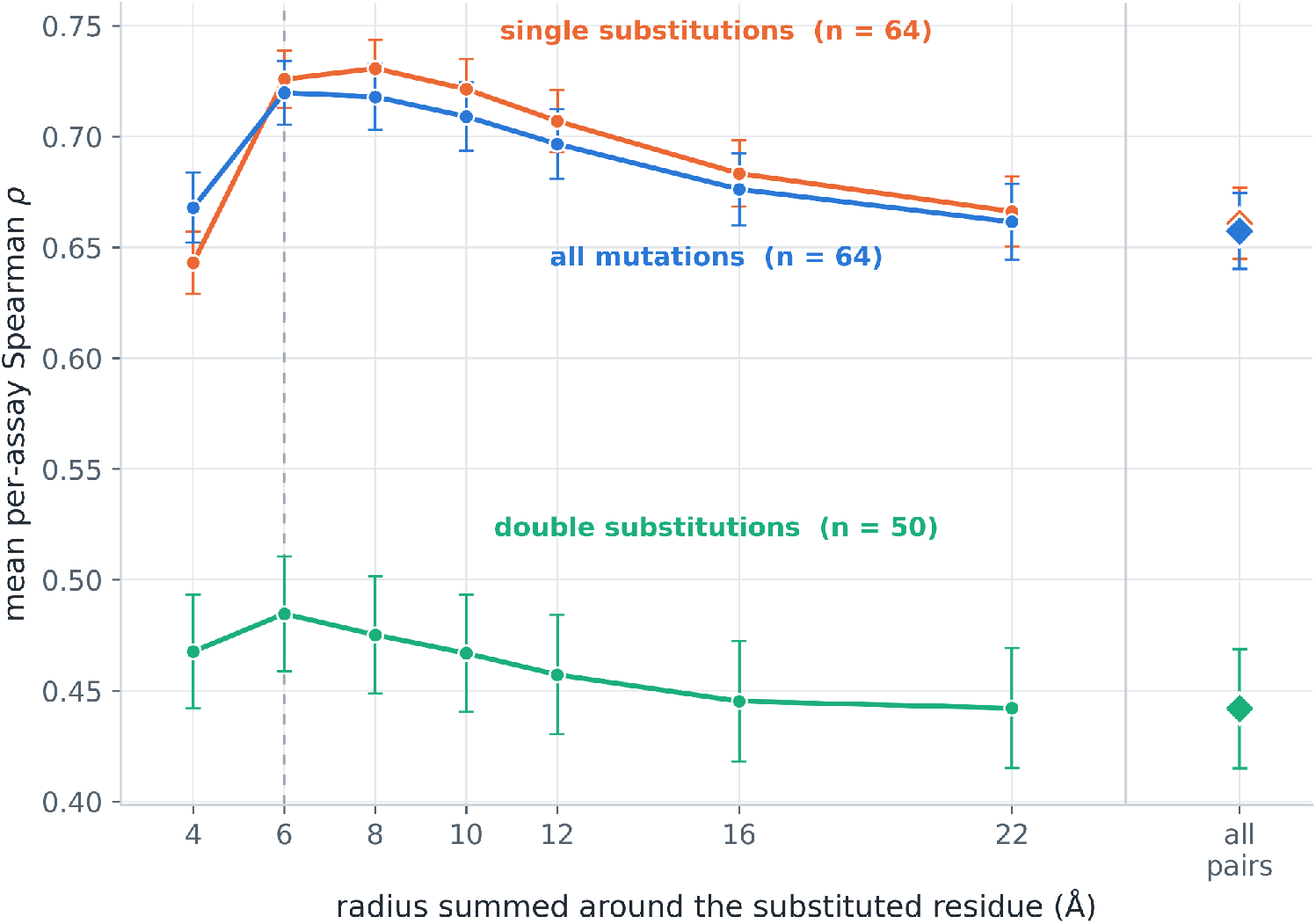
Spatial localization of mutation-sensitive energetic information. Mean per-assay Spearman correlation across 64 ProteinGym Stability assays as a function of the spatial cutoff around the mutation site. Predictive performance peaks at 6-8 Å and declines as more distant residue pairs are included.

To distinguish the physical locality of the signal from benchmark-specific optimization, the 6 Å formulation was selected using a predefined set of 16 development assays and subsequently evaluated without modification on the remaining 48 validation assays, which were not used for radius selection.

We further tested this spatial dependence on the independent S669 dataset and observed the same qualitative profile, with a broad optimum around 8 Å despite transferring the 6 Å formulation without retuning (Supplementary Note 2). The recurrence of this locality pattern in an independent dataset indicates that the benefit of restricting the score to the mutation neighbourhood is not specific to the ProteinGym proteins. Instead, the spatial decomposition reveals an intrinsic organization of the learned energetic response. Experimentally relevant mutation-sensitive information is concentrated in the immediate structural environment of the substituted residue, whereas progressively more distant responses contribute predominantly non-informative variation.

### 2.4 DEEP recovers the conformational landscape encoded by structure predictors

The mutation experiments establish that DEEP captures experimentally relevant energetic responses to sequence variation. We next asked whether the extracted potential also reflects the conformational preferences encoded by the underlying structure predictor. Specifically, without using RMSD, TM-score, or decoy quality during scoring, does DEEP assign progressively less favorable scores to increasingly non-native conformations?

We evaluated this property using the Rosetta decoy benchmark, comprising 180,079 structures across 133 protein targets ^33^. For each target, DEEP-Boltz scores were calculated across the corresponding conformational ensemble and compared with RMSD and TM-score relative to the native structure. Using sequence alignments, DEEP-Boltz reaches a mean per-target Spearman correlation of 0.884 with RMSD and 0.942 with negative TM-score. The native structure is the best-scoring conformation for 92% of targets, and the ten highest-scoring structures have a mean RMSD of 1.24 Å. Thus, the DEEP potential strongly orders alternative conformations according to structural nativeness across a large and diverse decoy collection.

This conformational signal does not depend entirely on evolutionary information. When Boltz-2 is run in single-sequence mode, DEEP retains mean correlations of 0.663 with RMSD and 0.710 with negative TM-score. For comparison, the Rosetta energy function reaches correlations of 0.750 and 0.759, respectively.

Because sequence is identical across all conformations of a target, the terminal separated-state reference is constant and cancels from within-target comparisons. Conformational discrimination therefore depends only on how the learned distance distributions score the geometries occupied by each structure.

Together, these results show that DEEP recovers both experimentally relevant mutation effects and the conformational preferences encoded by Boltz-2, raising the question of whether the same behavior is observed for another structure predictor.

### 2.5 DEEP generalizes to AlphaFold2 and separates structural recognition from mutation sensitivity

Having established both mutation sensitivity and conformational discrimination in DEEP-Boltz, we asked whether these properties necessarily emerge together across structure predictors. To address this question, we implemented the same DEEP framework using AlphaFold2 distograms (DEEP-AF2). DEEP requires a sequence-conditioned residue-pair distance distribution but is otherwise independent of the architecture of the underlying predictor. The same model-internal construction was therefore retained, including the terminal distance state as reference, allowing the energetic representations extracted from two independently developed structure predictors to be compared within a common framework.

The two predictors differed substantially when sequence was perturbed at fixed structure. Because AlphaFold2 inference is considerably more computationally demanding, we performed the cross-model comparison on a balanced subsample of 6,144 ProteinGym mutations, comprising 96 mutations from each of the 64 assays. Using the 6 Å local formulation and the same MSA depth of 256 for both predictors, DEEP-AF2 reaches a mean per-assay Spearman correlation of 0.583, compared with 0.723 for DEEP-Boltz. DEEP-Boltz performs better in 58 of the 64 assays, with a mean difference of +0.140 (Figure 4A). Despite this difference in predictive accuracy, the per-assay performances of the two models remain substantially correlated (ρ = 0.685, Figure 4C), indicating that they capture overlapping mutation-sensitive information but differ markedly in how quantitatively that information tracks experimental stability. The difference is robust to MSA depth, with AlphaFold2 showing little additional improvement beyond shallow alignments (Supplementary Note S4).

**Figure 4:**
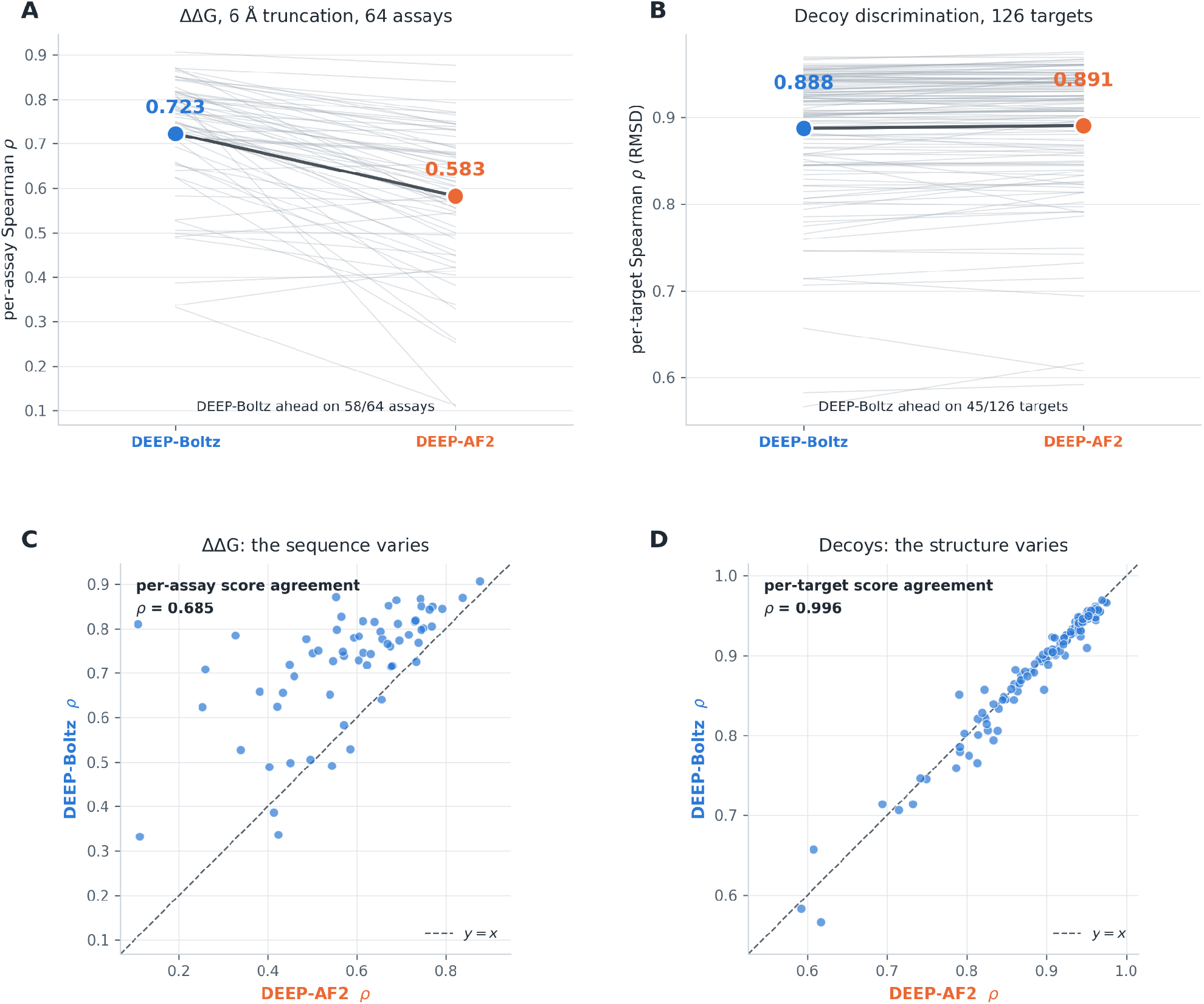
AlphaFold2 and Boltz-2 encode similar conformational preferences but differ in mutation-sensitive energetic response. (A) Mutation-effect prediction using DEEP-Boltz and DEEP-AF2 with the 6 Å local formulation across 6,144 ProteinGym mutations spanning 64 assays. Both models were evaluated using the same MSA depth of 256. Lines connect results for the same assay. (B) Conformational discrimination across 126 matched protein targets. In contrast to mutation prediction, DEEP-Boltz and DEEP-AF2 achieve nearly identical correlations. Lines connect results for the same target. (C) Per-assay mutation-prediction performance of the two models remains correlated (ρ = 0.685), indicating overlapping but quantitatively distinct responses to sequence perturbation. (D) Per-target conformational-discrimination performance is almost identical between the two models (ρ = 0.996). Together, these results show that conformational recognition and mutation-sensitive energetic response are separable properties of the learned representations.

Despite this difference in predictive accuracy, the spatial organization of mutation-sensitive information was remarkably similar between the two models. Repeating the spatial decomposition with DEEP-AF2 on the same 6,144 mutations revealed a maximum mean per-assay Spearman correlation of 0.583 at 6 Å, compared with 0.455 when all residue pairs were included (Supplementary Note S5). Local truncation improved performance in 62 of 64 assays, with a substantially larger benefit than observed for DEEP-Boltz. Thus, although the models differ in how accurately their mutation-sensitive responses predict experimental stability, the informative contributions are concentrated within a similar local structural environment.

A strikingly different picture emerges along the conformational axis. When sequence is held fixed and the same structures are scored across matched decoy ensembles, AlphaFold2 and Boltz-2 become nearly indistinguishable. Across 126 targets, DEEP-AF2 and DEEP-Boltz reach mean RMSD correlations of 0.891 and 0.888 (Figure 4B), respectively, and negative TM-score correlations of 0.948 and 0.947. More importantly, their per-target conformational discrimination scores agree almost perfectly (Figure 4D). Thus, the two models exhibit nearly identical conformational-discrimination performance despite substantially different mutation-sensitive responses.

The contrast between these two experiments reveals an important property of learned structural representations. Accurate recognition of the conformational landscape does not uniquely determine how it responds to changes in sequence. AlphaFold2 and Boltz-2 can agree almost perfectly on which conformations are compatible with a sequence while differing substantially in whether the energetic consequences of a mutation agree with experiment. Conformational recognition and mutation-sensitive energetic response are therefore related but separable learned properties of a structure predictor. The particularly strong mutation-sensitive signal in Boltz-2 consequently reflects information beyond that required simply to recognize native-like conformations.

### 2.6 DEEP captures mutation sensitive structural information complementary to existing predictors

The comparison between Boltz-2 and AlphaFold2 indicates that mutation-sensitive energetic information is not simply a consequence of learning to recognize native-like structures. We therefore asked whether the DEEP-Boltz signal also provides information that is distinct from sequence-based estimates of mutational compatibility. ODP2_GEOSE (PDB 1W4G) provides an informative case study because DEEP-Boltz and leading published predictors differ substantially across a densely sampled mutational landscape.

Across 1,134 experimentally characterized mutations in ODP2, DEEP-Boltz reaches a Spearman correlation of 0.545, compared with 0.390 for the strongest published method on this protein (Figure 5A). The difference is particularly informative at individual positions where evolutionary sequence preferences and experimentally measured stability diverge.

**Figure 5:**
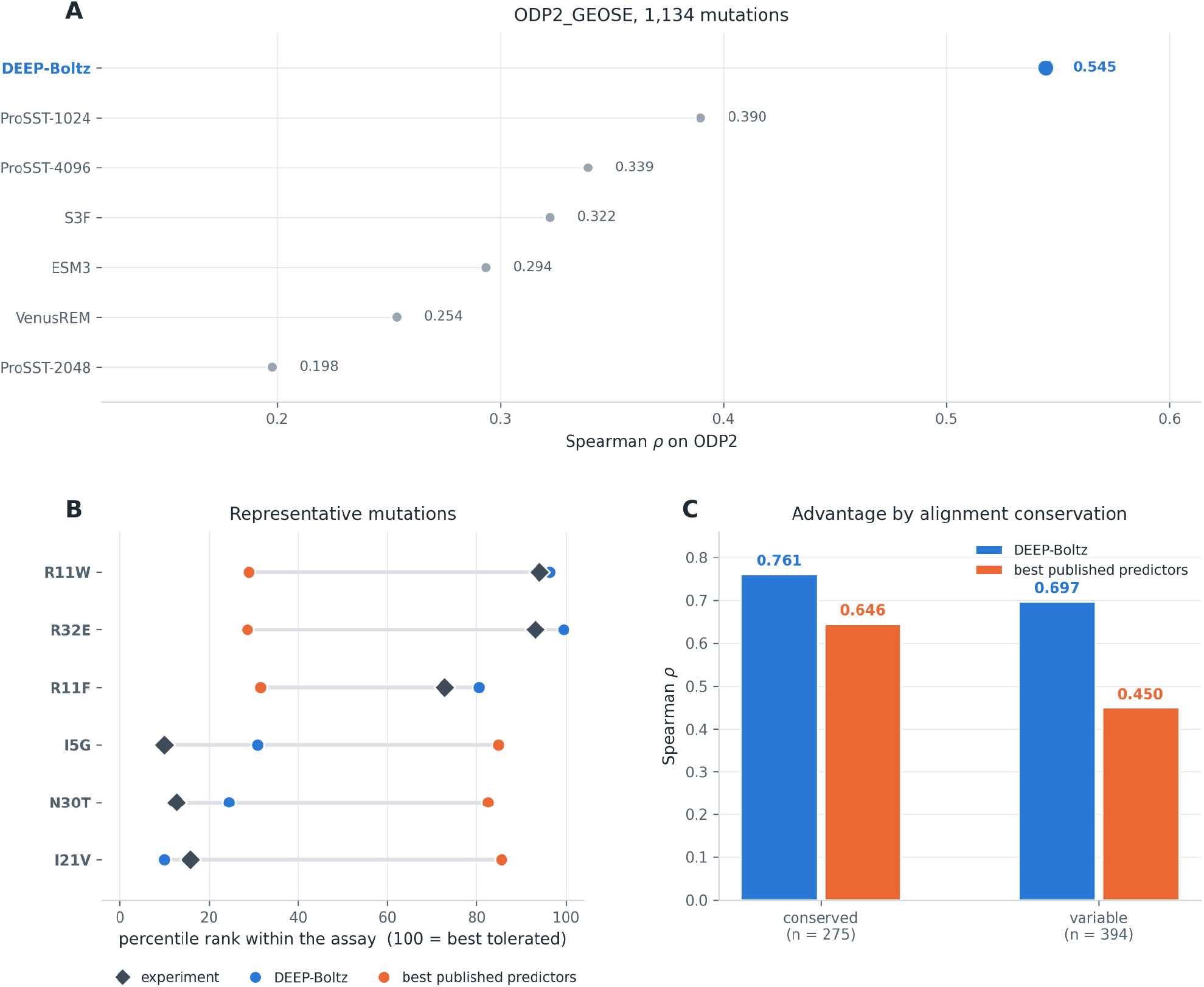
DEEP-Boltz captures mutation-sensitive structural constraints that complement sequence-based predictors in ODP2. **(A)** Mutation-effect prediction across all 1,134 experimentally characterized ODP2_GEOSE variants, compared with the leading published ProteinGym predictors. **(B)** Representative substitutions illustrating differences between experimental measurements, DEEP-Boltz and published predictions. Values are percentile ranks, with higher values indicating greater experimental tolerance or more favorable predicted effects. **(C)** Predictive performance by evolutionary conservation, evaluated on 669 single substitutions: 275 at conserved positions and 394 at variable positions. Positions were classified by a median split of alignment-derived Shannon entropy. The published method baseline represents the mean of the individual Spearman correlations of the six leading ProteinGym predictors within each group.

For example, R11W and R32E are radical substitutions that published mutation effect predictors strongly disfavor, yet both are experimentally well tolerated. DEEP-Boltz assigns these substitutions substantially more favorable scores, consistent with their measured effects. Conversely, I5G and N30T are experimentally poorly tolerated but are ranked among the more favorable substitutions by the published predictors; DEEP-Boltz assigns both substantially less favorable scores, closer to experiment. I21V provides a particularly informative structural example: although chemically conservative, the substitution removes a single methyl group from a buried isoleucine and produces an experimental stability loss of approximately 1.2 kcal/mol, which DEEP-Boltz correctly identifies as deleterious (Figure 5B). Together, these examples illustrate how an explicitly structural energetic representation can distinguish sequence compatibility from the energetic consequences of perturbing a particular three-dimensional environment.

To examine whether this complementarity depends on evolutionary conservation, we divided ODP2 positions into conserved and variable groups based on their alignment-derived Shannon entropy. The analysis included 669 single substitutions across 22 conserved and 22 variable positions. DEEP-Boltz achieved Spearman correlations of 0.761 at conserved positions and 0.697 at variable positions, compared with mean correlations of 0.646 and 0.450, respectively, across the six leading published predictors (Figure 5C). Thus, although predictive accuracy decreased at variable positions for both approaches, the decrease was substantially smaller for DEEP-Boltz. Its advantage over the published method baseline consequently increased from 0.115 at conserved positions to 0.247 at variable positions. This suggests that the structural information captured by DEEP remains predictive even where evolutionary sequence constraints are less informative.

The ODP2 landscape therefore illustrates the complementarity between sequence preference and structurally informed energetic response. Their disagreement at experimentally informative substitutions shows that the signal extracted by DEEP is not reducible to sequence preference alone.

## 3. Materials and Methods

### 3.1 Structure-predictor inference

DEEP-Boltz was implemented using sequence-conditioned distograms from the frozen Boltz-2 ‘boltz2_conf.ckpt’ checkpoint, loaded according to the Boltz 2.2.1 prediction pipeline. No parameters of the underlying model were trained, fine-tuned, or adapted ^2^. Boltz-2 inference used three recycling steps and full fp32 matrix multiplication.

For the primary DEEP-Boltz analyses, multiple-sequence alignments (MSAs) were limited to a depth of 256 sequences. ProteinGym alignments were converted from A2M to A3M format by uppercasing aligned residues and replacing ‘.’ characters with ‘-’, thereby preserving alignment register with the target sequence ^30^. Direct comparisons between Boltz-2 and AlphaFold2 used the same alignments truncated to a maximum depth of 256 sequences ^1^.

### 3.2 DEEP implementation

The DEEP statistical potential and mutation score are defined in Section 2.1. Unless otherwise stated, mutation analyses used the 6 Å local formulation.

For DEEP-Boltz, residue-pair distances were defined between Cβ atoms, with Cα used for glycine, following the residue-centre convention of Boltz-2. Cβ coordinates were reconstructed from backbone geometry using the same procedure as the Boltz-2 featurization pipeline. Self-pairs were excluded from all calculations.

### 3.3 Fixed-backbone mutation scoring

Mutation scoring followed the fixed-backbone formulation defined in Section 2.1. For each variant, the wild-type coordinates were retained and only the sequence-dependent model inputs were changed. Structural coordinates, pair weights, and the set of included residue pairs were therefore defined from the wild type and held fixed between wild-type and mutant calculations.

Substitutions to or from glycine require special treatment because Boltz-2 uses Cα rather than Cβ as the residue centre for glycine. Native distance-bin assignments were therefore recomputed for the mutant sequence where required, while retaining the wild-type pair weights and spatial mask. This ensures that wild-type and mutant scores are evaluated over the same set of residue pairs.

To assess sensitivity to the source of the reference structure, we repeated mutation scoring using wild-type structures generated by Boltz-2 instead of the AlphaFold2-derived reference structures distributed with ProteinGym. For each of the 64 assays, a single wild-type structure was generated using Boltz-2’s structure module, conditioned on the same trunk representation used for mutation scoring. Structures were sampled using 200 reverse-diffusion steps, the full-atom input layout, and an assay-specific deterministic random seed. Each predicted backbone was subsequently held fixed across all variants of the corresponding assay.

The comparison included all 117,811 variants, using identical mutant trunk outputs, alignments (maximum depth 256), model inference settings, and scoring parameters. Pair weights were recalculated for each reference structure using the same weighting procedure. The two conditions therefore differed only in the structural coordinates used to evaluate the mutation scores. This comparison was initially performed using the complete, untruncated residue-pair sum, rather than the primary 6 Å local formulation. Performance differences were assessed using paired per-assay Spearman correlations, with confidence intervals obtained by bootstrap resampling and significance evaluated using a Wilcoxon signed-rank test.

### 3.4 ProteinGym stability benchmark

Mutation-effect prediction was evaluated using 64 ProteinGym v1.3 Stability assays comprising 117,811 experimentally characterized variants, including 65,887 single and 51,924 double substitutions ^30^. One reference structure was used for each assay. Three subsets of this benchmark were used for distinct purposes.

The primary ProteinGym evaluation used the complete set of 117,811 variants across all 64 assays. This dataset was used to report the main DEEP-Boltz performance and to compare against published ProteinGym predictors.

To evaluate whether methodological choices generalized beyond the assays used to select them, 16 development assays were selected deterministically by sorting the 64 assays by wild-type sequence length and taking evenly spaced entries across the resulting range. The remaining 48 assays were used for validation. The 6 Å spatial cutoff was selected using only the 16 development assays and subsequently evaluated, without modification, across the full mutation sets of the 48 validation assays. The same development/validation partition was used for the weighting and MSA-depth analyses.

Finally, computationally intensive analyses requiring repeated AlphaFold2 inference used a deterministic balanced subsample of 6,144 variants. All 64 assays were represented by 96 variants each. Within each assay, variants were sorted by experimental DMS score and 96 evenly spaced entries were selected across the resulting rank distribution, ensuring coverage of the measured mutational range without random sampling. The same 6,144 variants were used for DEEP-Boltz and DEEP-AF2 in all direct cross-model comparisons and AlphaFold2 MSA-depth analyses. This balanced subsample is distinct from the assay-level development/validation split described above.

Performance was calculated separately within each assay using Spearman rank correlation between the DEEP mutation score and the experimental stability measurements. Overall performance was summarized as the mean per-assay Spearman correlation. Published ProteinGym per-assay scores were re-aggregated over the same 64 assays to provide matched comparisons with other zero-shot mutation-effect predictors.

### 3.5 Independent S669 evaluation

Independent validation was performed using the S669 protein-stability benchmark, which contains 669 experimentally characterized single substitutions across 94 protein chains. Six chains with reported homology to the Tsuboyama/Megascale collection were excluded from the independent analysis ^32,34^. Following sequence-structure reconciliation and technical exclusions, the independent evaluation comprised 488 variants.

Because S669 contains relatively few variants for many proteins and combines measurements from heterogeneous experimental sources, performance was evaluated both within individual proteins and across the pooled dataset. Within-protein performance was calculated for proteins containing at least five variants and summarized as the mean Spearman correlation across proteins. Pooled Spearman and Pearson correlations were calculated separately. Comparisons with published S669 methods were restricted to variants for which the corresponding baseline predictions were available. Detailed filtering, baseline matching, and analyses of the alternative evaluation protocols are provided in Supplementary Note S2.

### 3.6 Conformational discrimination

Conformational discrimination was evaluated using the 133-target Rosetta decoy benchmark, comprising 180,079 alternative protein conformations ^33,35^. Each target consists of an experimentally determined native structure and a collection of decoys sharing the same amino-acid sequence.

Each conformation was scored using the complete DEEP potential. For conformational scoring, the radial volume correction defined in Section 2.1 was applied. Resolved distance bins were treated as spherical shells, whereas the terminal distance state was represented by a sphere truncated at 45 Å.

Within each target, DEEP scores were compared with RMSD and negative TM-score relative to the native structure using Spearman rank correlation ^36^. Performance was summarized as the mean per-target correlation. Native-state recognition was additionally evaluated from the rank assigned to the experimentally determined structure, and the structural quality of the highest-scoring conformations was evaluated from their RMSD.

MSAs for the decoy targets were generated using the ColabFold MMseqs2 server ^37,38^. Alignments were successfully obtained for 126 of the 133 targets; these 126 targets constituted the matched set used for direct comparison between DEEP-Boltz and DEEP-AF2. Single-sequence calculations were performed by running the same scoring procedure without evolutionary sequence information.

### 3.7 AlphaFold2 implementation

DEEP-AF2 was implemented from the sequence-conditioned AlphaFold2 distogram output using the same general formulation as DEEP-Boltz. AlphaFold2 represents residue-pair distances using 64 bins spanning approximately 2.31-21.69 Å; its own terminal distance bin was used as the corresponding model-internal far-distance state.

Because AlphaFold2 inference was substantially more computationally demanding, mutation analysis was performed on the deterministic balanced 6,144-variant ProteinGym subsample defined in Section 3.4. For direct cross-model comparisons, DEEP-Boltz and DEEP-AF2 were evaluated on identical variants using the same MSAs truncated to a maximum depth of 256 sequences and the same 6 Å spatial cutoff.

To assess sensitivity to MSA depth, DEEP-AF2 was additionally evaluated at maximum alignment depths of 1, 32, 64, 128, 512, and 1024 sequences while keeping all other inference and scoring settings fixed. The depth-1 condition retained only the query sequence and therefore served as the single-sequence control. All depth comparisons were performed on the same 6,144 mutations.

Conformational discrimination was compared between the two models on the 126 Rosetta targets for which alignments were available. Identical conformations and sequence alignments were supplied to both models, and each predictor’s own terminal distance state was used as its DEEP reference.

### 3.8 Evolutionary conservation analysis

Evolutionary conservation of ODP2_GEOSE was calculated from the ProteinGym v1.3 multiple-sequence alignment (ODP2_GEOSE_2023-08-07_b07), using the first 4,000 of 163,835 sequences and restricting the analysis to columns corresponding to the 44-residue query sequence. Conservation was quantified by positional Shannon entropy, with lower entropy indicating greater conservation ^39^. Positions were classified using the median entropy across the 44 residues (1.43 bits), yielding 22 conserved positions (entropy ≤ 1.43 bits) and 22 variable positions (entropy > 1.43 bits). Calculating the median across positions rather than variants prevents differences in mutational sampling density from influencing the classification.

The analysis was restricted to single substitutions at positions with defined entropy, yielding 669 of the 1,134 experimentally characterized ODP2 variants, comprising 275 substitutions at conserved positions and 394 at variable positions. Predictive performance was evaluated separately within each group using Spearman rank correlation. DEEP-Boltz was compared with the six highest-performing published ProteinGym predictors on ODP2 (ProSST-K1024, ProSST-K2048, ProSST-K4096, VenusREM, ESM3 and S3F) ^40-43^. The published-method baseline was calculated as the arithmetic mean of the six individual Spearman correlations within each conservation group, rather than the correlation of averaged prediction scores. All methods were evaluated on identical variants within each group.

## 4. Discussion

Modern protein structure predictors learn representations that extend beyond the coordinates they generate, but the energetic information contained within these representations remains incompletely understood. Here, we demonstrate that sequence-conditioned distograms can be transformed into effective statistical potentials that capture both experimentally measured mutation effects and conformational preferences. DEEP extracts this information directly from frozen structure predictors without training on experimental stability data. Its strong performance in zero-shot stability prediction and conformational discrimination establishes that learned structural representations contain accessible energetic information with practical predictive value.

A central feature of DEEP is its use of the predictor’s own terminal distance state as an internal reference. Unlike classical statistical potentials, which rely on externally defined background distributions, DEEP evaluates an observed residue-pair geometry relative to a model-encoded state of spatial separation. The effectiveness of this reference across Boltz-2 and AlphaFold2 suggests that the terminal state contains information beyond that represented by ordinary finite-distance bins. Nevertheless, this state should not be interpreted as a thermodynamic unfolded ensemble, nor should the resulting potential be regarded as a physically calibrated free energy. Rather, DEEP provides an operational energetic representation of the sequence-structure preferences learned by the predictor. The strong localization of mutation-sensitive information within approximately 6-8 Å of the substituted residue further suggests that this representation captures changes in the compatibility of the immediate structural environment rather than simply assigning intrinsic preferences to individual amino acids.

Perhaps the most important conceptual finding emerges from comparing Boltz-2 and AlphaFold2. Despite nearly identical conformational discrimination, the two predictors differ substantially in their ability to capture experimentally measured mutation effects. Thus, accurately recognizing native-like conformations does not necessarily imply that a model has learned how structural preferences should respond to changes in sequence. Conformational recognition and mutation-sensitive energetic response are therefore related but distinct properties of learned structural representations. This distinction suggests that evaluating structure predictors solely by geometric accuracy overlooks an important dimension of their learned behavior. DEEP offers a complementary means of interrogating these representations by examining both their conformational preferences and their sensitivity to sequence perturbations.

Beyond these conceptual implications, DEEP provides a practical approach to mutation-effect prediction. DEEP-Boltz achieves leading zero-shot performance on ProteinGym and remains competitive on the independent S669 benchmark without fitting to experimental stability measurements. The ODP2 case study further demonstrates that its predictions can complement sequence-based approaches, particularly where evolutionary preferences do not fully reflect the energetic consequences of substitutions in a specific structural environment. These findings suggest that structure-derived energetic representations provide information that is not entirely captured by sequence-based mutation predictors.

Several limitations remain. DEEP scores are effective statistical potentials rather than absolute thermodynamic free energies, and their scales may differ between proteins, limiting direct cross-protein comparisons. The fixed-backbone mutation formulation does not explicitly account for structural relaxation, changes in conformational ensembles, or large-scale conformational transitions. Moreover, although the reference-state and spatial analyses identify informative components of the learned representations, they do not establish the mechanisms by which different model architectures acquire this information. Addressing these limitations may improve quantitative stability prediction and clarify the relationship between learned structural probabilities and physical energetics. More broadly, DEEP establishes a framework for evaluating protein structure predictors not only by the structures they generate, but also by the energetic information encoded in their internal representations.

## Code availability

The DEEP framework and its DEEP-Boltz implementation are freely available as open-source software at https://github.com/erdoslab-elte/DEEP-Boltz. The repository provides the implementation and instructions for extracting statistical potentials from protein structure predictors and predicting mutation induced protein stability changes.

## Funding

This work was funded by the European Union’s Horizon Europe programme under grant agreement No. 101160233 (IDP2Biomed, HORIZON-WIDERA-2023) and grant agreement No. 101182949 (IDPfun2, HORIZON-MSCA-2023-SE). E.G. acknowledges funding from EKA: 2025/041-P108-1, ELIXIR CMR 2024/2025 and FEBS Booster Funds 8239097.

## Disclaimer

“Views and opinions expressed are however those of the author(s) only and do not necessarily reflect those of the European Union or Research Executive Agency. Neither the European Union nor the granting authority can be held responsible for them.”

## Acknowledgements

The authors thank Zsuzsanna Dosztányi and Richárd Hornyák for helpful conversations. Claude Code (Opus 4.5 / Sonnet 4.5) was employed in the implementation of this work.

## Conflict of interest statement

None declared.

## Notes

https://github.com/erdoslab-elte/DEEP-Boltz

### Competing Interest Statement

The authors have declared no competing interest.

### Summary of Updates

Updated method section and supplementary information

