## Supplementary Information for "DEEP: Extracting Latent Energetic Landscapes from Protein Structure Predictors"

#### Supplementary Note S1. Reference state and far bin decomposition

We evaluated 67 reference definitions on the balanced 6,144-variant ProteinGym subsample described in Section 3.4, comprising 96 variants from each of the 64 assays. Under the primary 6 Å formulation, the terminal distance state produced the highest mean per-assay Spearman correlation for both DEEP-Boltz and DEEP-AF2 (Supplementary Figure S1). For DEEP-Boltz, performance increased from 0.6086 without reference subtraction to 0.7148 using the terminal state. For DEEP-AF2, the corresponding increase was from 0.5435 to 0.5888.

Importantly, the advantage of the terminal state was not simply a continuation of a trend across progressively larger distance bins. For DEEP-Boltz, reference terms derived from the 63 finite-distance bins reached a mean correlation of 0.6388 ± 0.0149, compared with 0.7133 for the terminal state alone. For DEEP-AF2, the corresponding values were 0.4617 ± 0.0066 and 0.4993. The terminal state therefore behaves as a distinct component of the learned representation rather than merely as the outermost point of the resolved distance distribution.

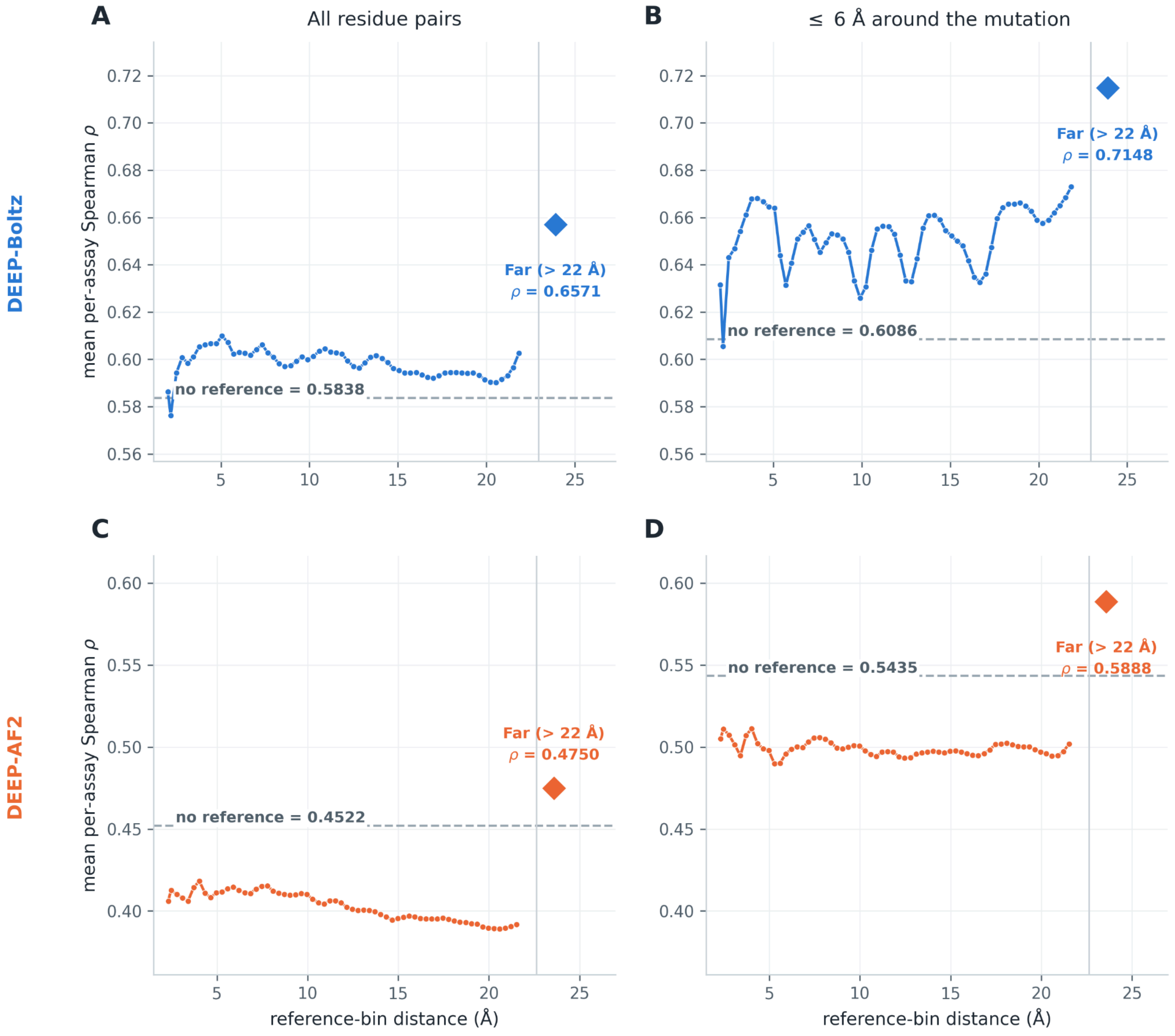


***Supplementary Figure S1: The terminal distance state provides a distinct and informative reference for DEEP.*** ***(A,B)*** *Mean per-assay Spearman correlation of DEEP-Boltz using individual distance bins as reference states with the complete pair set (A) and the 6 Å local formulation (B).* ***(C,D)*** *Corresponding analysis for DEEP-AF2. Dashed lines indicate performance without reference subtraction; diamonds indicate the terminal open-ended distance state. The terminal state outperforms all finite-distance reference bins under the primary 6 Å formulation in both models.*

### Supplementary Note S2: S669 benchmark set validation

We evaluated DEEP-Boltz on the independent S669 benchmark to assess the generalizability of mutation-effect prediction and the spatial localization observed on ProteinGym. S669 comprises 669 experimentally characterized single substitutions across 94 protein chains, collected from 93 publications under heterogeneous experimental conditions. To minimize overlap with the Tsuboyama/Megascale dataset, six chains with reported homologues in that collection were excluded. Following sequence–structure reconciliation and technical filtering, the independent evaluation comprised 488 variants.

Because S669 contains relatively few substitutions per protein and combines measurements from different experimental sources, we primarily assessed the ability of DEEP-Boltz to rank mutation effects within individual proteins. Among the 21 proteins containing at least five variants, DEEP-Boltz achieved a mean within-protein Spearman correlation of 0.413 using the fixed 6 Å formulation established on ProteinGym. This ranked DEEP-Boltz third among 21 methods evaluated on matched variants, behind INPS3D (0.461) and PremPS (0.415), and first among the zero-shot predictors. These results demonstrate that DEEP-Boltz retains competitive mutation-ranking performance on an independent, experimentally heterogeneous stability benchmark without retraining or modifying its scoring parameters.

We further examined whether the spatial localization of mutation-sensitive information generalized to S669 by evaluating the dependence of predictive performance on the structural cutoff around the substituted residue. Consistent with ProteinGym, prediction accuracy exhibited a broad optimum around 8 Å, with more distant residue-pair contributions reducing performance. Importantly, the primary 6 Å formulation was transferred without retuning, supporting the generalizability of the localized mutation-sensitive signal across independent datasets.

#### Supplementary Note S3. Ablation of residue-pair weighting

DEEP weights each residue-pair contribution by both the probability assigned to the observed geometry and the confidence of the corresponding distance distribution,

$$w_{ij} = \exp(-H_{ij})\, p_{ij}\left(\mathrm{bin}(d_{ij}) \mid s_{\mathrm{WT}}\right)$$

where $H_{ij}$ is the entropy of the predicted distogram. To determine the contribution of these two components, we compared the complete weighting with uniform weighting, observed-distance probability alone, and entropy-based confidence alone.

Across the complete set of 64 ProteinGym assays, uniform weighting reaches a mean per-assay Spearman correlation of 0.677, compared with 0.715 for observed-distance probability alone and 0.707 for entropy weighting alone. Combining the two terms gives the highest overall performance (0.720). On the predefined 48-assay validation set, weighting by observed-distance probability improves the mean correlation by 0.033 relative to uniform weighting, with improvements in 42 of 48 assays. Thus, most of the benefit of residue-pair weighting arises from emphasizing pairs for which the predictor assigns high probability to the geometry observed in the wild-type structure.

We further evaluated the incremental contribution of the entropy term using the complete production pipeline. For single substitutions, adding entropy weighting to observed-distance probability has essentially no effect: mean per-assay correlation changes from 0.7255 to 0.7258. The same is observed on the 48-assay validation set, where the entropy term slightly decreases mean performance. In contrast, for double substitutions the entropy term increases mean correlation from 0.4735 to 0.4847. This improvement is retained on the validation assays.

Accordingly, observed-distance probability provides the dominant weighting signal and is sufficient for essentially all of the performance on single substitutions, whereas entropy-based confidence provides a modest additional benefit for variants containing two substitutions. We therefore retain the combined probability and entropy weighting in the general DEEP formulation, which is applicable to both single and multiple substitutions. Importantly, the near-identical performance obtained for single substitutions with and without entropy weighting shows that the principal mutation-sensitive signal extracted by DEEP does not depend on the entropy-based confidence term.

#### Supplementary Note S4. Mutation prediction is robust to MSA depth

Because DEEP extracts sequence-dependent information from structure-predictor distograms, we tested how strongly mutation prediction depends on the amount of evolutionary information supplied to AlphaFold2. DEEP-AF2 was evaluated on the same 6,144 ProteinGym mutations at maximum MSA depths of 1, 32, 64, 128, 256, 512, and 1024 sequences, with all other inference and scoring settings held fixed.

Removing alignment information substantially reduced performance: mean per-assay Spearman correlation decreased from 0.589 at depth 1024 to 0.406 when only the query sequence was supplied ($\Delta\rho = -0.184$, $p = 3.7 \times 10^{-8}$). In contrast, most of the alignment-dependent signal was already recovered with a shallow MSA. A depth of 32 sequences reached a mean correlation of 0.562, recovering approximately 85% of the improvement between single-sequence and depth-1024 inference. Increasing alignment depth further produced no consistent improvement: correlations were 0.566, 0.586, 0.583, 0.551, and 0.589 at depths 64, 128, 256, 512, and 1024, respectively, and none differed significantly from the depth-1024 condition.

Thus, evolutionary information substantially strengthens the mutation-sensitive signal extracted by DEEP-AF2, but this signal saturates at relatively shallow alignment depth. Importantly, the difference between DEEP-Boltz and DEEP-AF2 is not explained by unequal MSA depth. When both predictors are evaluated at the production depth of 256 sequences, DEEP-Boltz reaches 0.723 compared with 0.583 for DEEP-AF2, increasing the cross-model difference to 0.140.

#### Supplementary Note S5. Spatial localization of mutation-sensitive information in DEEP-AF2

To determine whether the spatial localization of mutation-sensitive information observed in DEEP-Boltz generalizes to AlphaFold2, we evaluated DEEP-AF2 across structural cutoffs ranging from 4 to 22 Å, together with the complete untruncated pairwise sum. The analysis used the balanced ProteinGym subset of 6,144 mutations across 64 assays, with a maximum MSA depth of 256 sequences.

DEEP-AF2 exhibited a pronounced local optimum at 6 Å, reaching a mean per-assay Spearman correlation of 0.583, compared with 0.455 when all residue pairs were included (Supplementary Figure S2). Restricting the score to 6 Å improved performance in 62 of 64 assays. The improvement of +0.128 was substantially greater than the corresponding improvement observed for DEEP-Boltz, indicating that distant pairwise contributions reduce mutation-prediction accuracy more strongly in DEEP-AF2.

The spatial dependence was robust to MSA depth, with an independent evaluation at depth 1024 recovering the same 6 Å optimum ($\rho = 0.589$). Single substitutions exhibited a maximum at 6 Å ($\rho = 0.572$), whereas double substitutions peaked at 4 Å ($\rho = 0.458$). Thus, despite differences in absolute predictive accuracy and some variation between mutation classes, both structure predictors concentrate experimentally informative mutation-sensitive responses within a similar local structural environment.

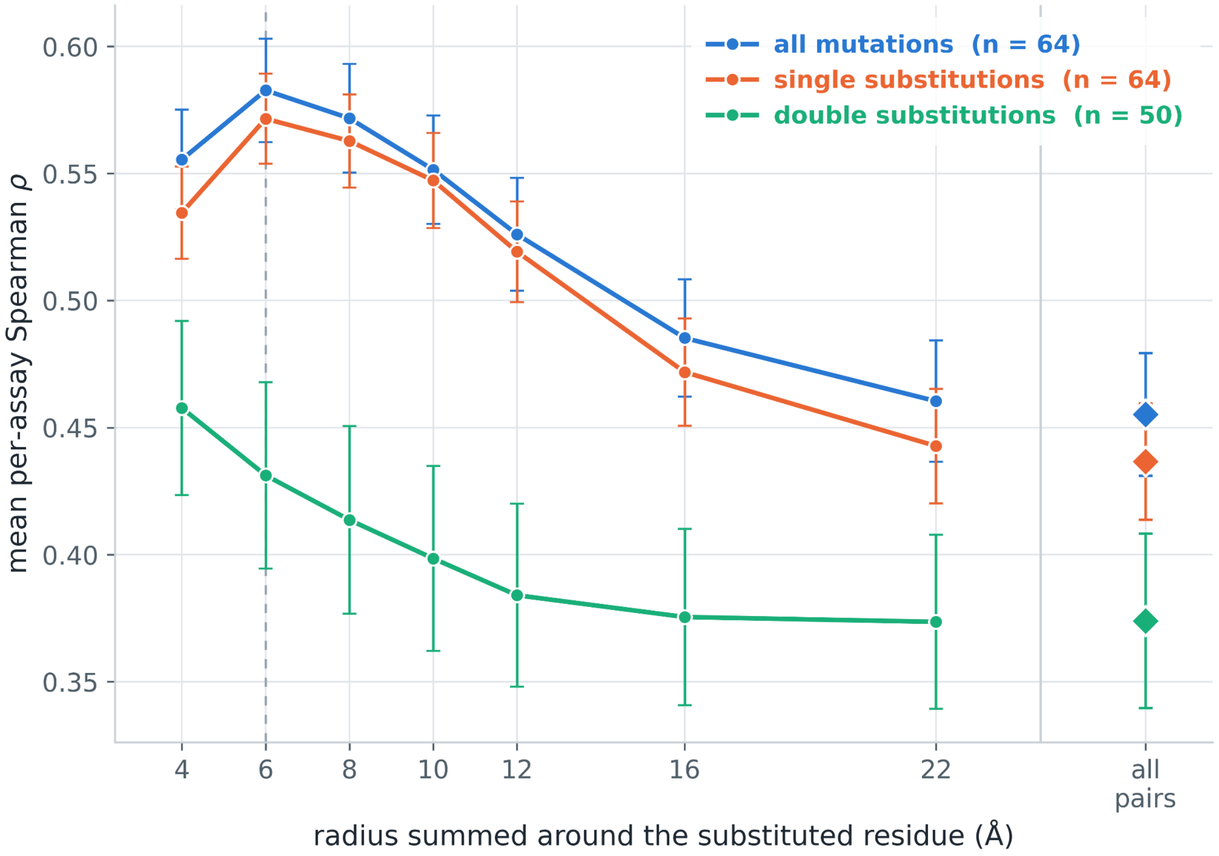


***Supplementary Figure S2. Spatial localization of mutation-sensitive information in DEEP-AF2.*** *Mean per-assay Spearman correlation as a function of structural cutoff across 6,144 ProteinGym mutations from 64 assays. DEEP-AF2 achieves maximum predictive performance at 6 Å, with a substantially greater benefit from spatial truncation than DEEP-Boltz. The local optimum is preserved at MSA depths of 256 and 1024. Single substitutions peak at 6 Å, whereas double substitutions peak at 4 Å.*
